# Density dependence affects population recovery following an epidemic

**DOI:** 10.64898/2025.12.12.693954

**Authors:** Rustom Antia, Bert Baumgaertner, James J Bull

## Abstract

Current epidemiological models for the regulation of host populations by pathogens suggest that the reduction in the host population is largest shortly after the epidemic phase and that the host population partly recovers in the subsequent approach to endemicity. We find that this conventional view is not robust to a simple, biologically motivated addition: density-dependent population regulation (homeostasis). While rapid homeostasis typically enables a population rebound after the initial population suppression (the conventional view), we show that weak homeostasis can result in the opposite — greater population suppression during the endemic phase compared to the initial epidemic. The effect of homeostasis, which is a property of the host population, can be substantial. This finding complicates the prediction of long term outcomes of population suppression by pathogens but not of short term outcomes.

## Introduction

Viruses and other pathogens can cause substantial mortality when introduced into naive animal or human populations [1, 7, 13, 20, 25], and there is a long history of using epidemiological models to understand their effects in regulating and suppressing host populations [1, 12, 15]. Most of these modeling studies focus on how pathogens suppress host populations at steady-state when the pathogen and host numbers have equilibrated, which corresponds to the endemic phase. Other epidemiological models consider the spread of infections during the initial epidemic wave that corresponds to the first wave of infections following the introduction of a pathogen into a naive population [3, 17, 21, 22]. However these studies do not typically consider the suppression of the host population during this initial epidemic phase. Our focus here is to extend earlier studies to consider the effect of a pathogen on suppressing the host population in the initial epidemic phase and compare this with the extent of suppression after the pathogen has reached an endemic equilibrium. In particular we ask whether we see the largest population depression during the initial epidemic phase shortly after the first introduction or later in the approach to endemicity.

The first wave of infections in characterized by the explosive growth in the number of infecteds which reaches its peak when the number of susceptibles equals 1*/R*_0_. Following this peak the effective reproductive number of the pathogen is less than one, but the momentum of the epidemic results in a larger, and sometimes considerably larger fraction of the population becoming infected. The fraction of the population that becomes infected during the initial wave is called the “final size” of the epidemic phase, and the extent to which the number of susceptibles falls below 1*/R*_0_ is termed the “overshoot”. Approximations for the final size have been extensively studied since first described by Kermack and Mckendrick almost a hundred years ago [3, 17, 21, 22]. The overshoot in the number of infecteds and its relationship to the final size of an epidemic has also been discussed, particularly during the SARS-CoV-2 pandemic [14, 18, 22]. The final size and overshoot calculations are not affected by pathogen lethality, and this allows us to extend earlier analytical results to consider how the suppression of the host population during the epidemic phase depends on both the transmissibility (*R*_0_) and severity (case mortality) of infections.

On first consideration, one might expect the overshoot to result in higher population suppression during the initial epidemic phase than at endemism. In this note we use simple epidemic models to address whether this is the case. We compare short and long-term suppression of the host population by a pathogen. We do so by extending the final size equation to consider the degree of suppression of the host population in the short term. We then examine how the degree of suppression changes in the transition to endemicity.

## Models and Results

The models we consider in this paper are simple epidemiological models that consider three compartments of individuals: **S**usceptible, **I**infectious and **R**ecovered. We further assume that the infections are of short duration (so called acute infections) that result in lifelong immunity in the individuals who survive infection (i.e., who recover). The relevant equations are below, with notation and parameters in Table 1:

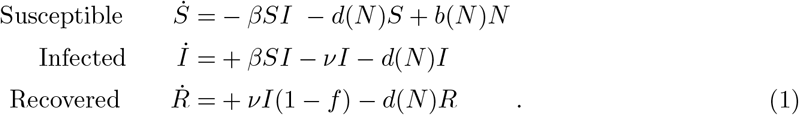

**Table 1:** Notation for variables and parameters in eqns (1). We analyze two different sets of values for the four birth-death parameters. Birth and death parameter values are chosen so that the equilibrium population size in the absence of the pathogen is 1, and the parameters for rapid homeostasis are chosen to represent a short-lived, high-fecundity species such as rabbits (see Supplement S1 for details); values for slow homeostasis are chosen to contrast with those for rapid homeostasis.

| Symbol | Description | Value |
| --- | --- | --- |
| $S$ | Susceptible, born of naive mothers | densities, $\leq 1$ |
| $I$ | Infected $S$ | |
| $R$ | Recovered, immune | |
| $N$ | Total population ($S + I + R$) | |
| $\nu$ | loss of infection rate | 50 |
| $\beta$ | transmission rate constants | for $R_0$ ($1.5 < R_0 < 10$) |
| $f$ | case mortality | $0 < f < 1$ per infection |
| Rapid homeostasis |  |  |
| $b_0, d_0$ | intrinsic birth and death rates | 3.5, 0.2 /year |
| $k_b, k_d$ | parameters for logistic growth | 2.5, 0.8 |
| Slow homeostasis |  |  |
| $b_0, d_0$ | intrinsic birth and death rates | 1.5, 0.8 /year |
| $k_b, k_d$ | parameters for logistic growth | 0.5, 0.2 |

*N* is total population size, here scaled to 1 for an equilibrium population lacking the infection. The severity of infections is described by the case mortality (*f*). We choose case mortality rather than a virulence rate (e.g. [11]) for two reasons. First it gives a more intuitive description of virulence than a rate of virulence, and second it facilitates exploring the effect of independently changing transmission and disease severity. The latter is in contrast with traditional models where virulence (*α*) is introduced by a term −*αI* in the equation for *dI/dt* – which results in changes in *α* also changing *R*_0_.

In our framework the basic reproductive number (*R*_0_) equals *β/*(*ν* + *d*(*N*)), or ≈ *β/ν*, an approximation that relies on the duration of infection being much shorter compared to host lifespan in a disease-free population. The simulations that vary *R*_0_ do so by changing transmission, *β*.

### Short-term Population Suppression: the final size equation

Following introduction of the pathogen into a naive host population, the number of infected individuals increases exponentially at first, slowing down as the number of susceptible individuals declines. There is a well-defined (approximate) endpoint to this initial phase, and the cumulate number/density of individuals infected is known as the ‘final size.’ These initial dynamics during the first wave occur on a timescale that are fast compared with the birth and death rates of hosts and is approximated by setting births and deaths to zero, enabling analytical solutions to the final size equation.

We can use the final size formula [3, 17, 21] to estimate the fraction of the population that has been infected (*F*) from the solution to

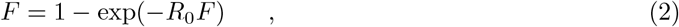

with *R*_0_ ≈ *β/ν*. The susceptibles remaining at the end of this initial phase will equal 1 − *F* (given our constraint that initial density is 1), and the number of individuals who have survived to recover is (1 − *f*)*F*, where *f* is the case mortality. This gives the minimum population during the epidemic phase as

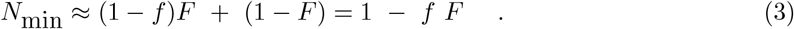

#### Excess deaths and the expectation of a population rebound

Our question is how the population will change beyond this initial phase: will it increase or decrease in the approach to endemicity? Intuitively, we might derive insight from a phenomenon known as ‘overshoot,’ which operates during the late phase of the epidemic. The overshoot of the epidemic equals the proportion of the population that becomes infected between the epidemic’s peak and the epidemic’s final size. The peak of the epidemic occurs when *S* = 1*/R*_0_, so the number of excess deaths will be:

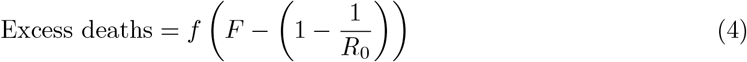

We might naively expect the number of excess deaths to give some indication of how the population will change toward the endemic phase. Since the excess death number is positive, we would conclude that the population always rebounds, i.e., that the epidemic final size is always smaller than the endemic size. However compelling that this inference may appear, it is wrong.

### Long-term population size and the effect of “homeostasis”

We have shown how the suppression of the host population during the initial or epidemic phase depends on its transmissibility (characterized by *R*_0_) and virulence (characterized by case mortality *f*). We now explore how long-term population suppression due to a pathogen depends not only on *R*_0_ and *f* but also on the intrinsic regulation of the host population.

To simplify the terminology and to emphasize the compensatory impact of density dependence, we refer to density dependence as the “homeostatic” regulation of the host population. Homeostasis can be rapid or slow, depending on the assumptions and parameterization of the model. In model (1), population regulation stems from the generic terms *b*(*N*) and *d*(*N*) that describe the birth and death rates for the host population.

Various models can describe homeostasis in the absence of the pathogen. We analyze two models for homeostatic regulation of the host population. The first, and simpler model assumes a constant number of births per unit time, and where individuals the death rate is fixed (i.e. *d*(*N*) = *d*). In the second model we assume logistic growth where the birth rate declines linearly and the death rate increases linearly with population size (i.e. *b*(*N*) = *b*_0_ − *k*_*b*_*N* and *d*(*N*) = *d*_0_ + *k*_*d*_*N*).

#### Constant births and density dependent deaths

In this model, the equations governing the dynamics of pathogen and host are:

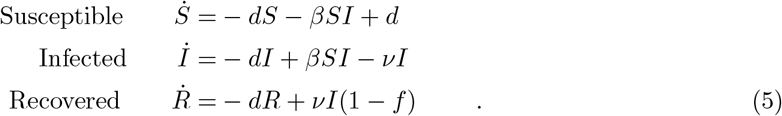

Without loss of generality, the number of births is set equal to the death rate so the pathogen-free population density equals 1 at steady-state (*Ŝ* = 1). Homeostasis is due entirely to deaths being less than births when the total population size *N* < 1 and greater than births when *N* > 1. With the pathogen present, the endemic equilibrium is:

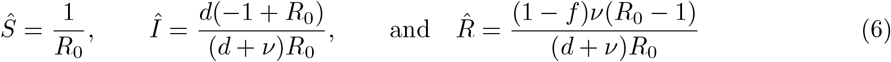

The total population size at the endemic steady-state, 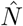 equals:

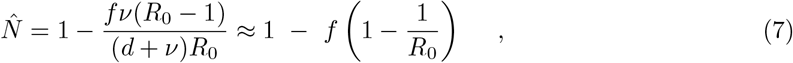

where the approximation relies on the duration of infection being much smaller than the lifespan (i.e. *ν* >> *d*).

Fig 1 panel A plots the minimum host population reached during the initial epidemic phase (calculated from eqn (3)) and panel B plots the host population size at endemicity (calculated from the exact solution in eqn (7)). Suppression during the epidemic phase is slightly higher than during the endemic phase. This is the conventional view of the relationship between epidemic and endemic phases and agrees qualitatively with the inference based on overshoot.

**Figure 1:**
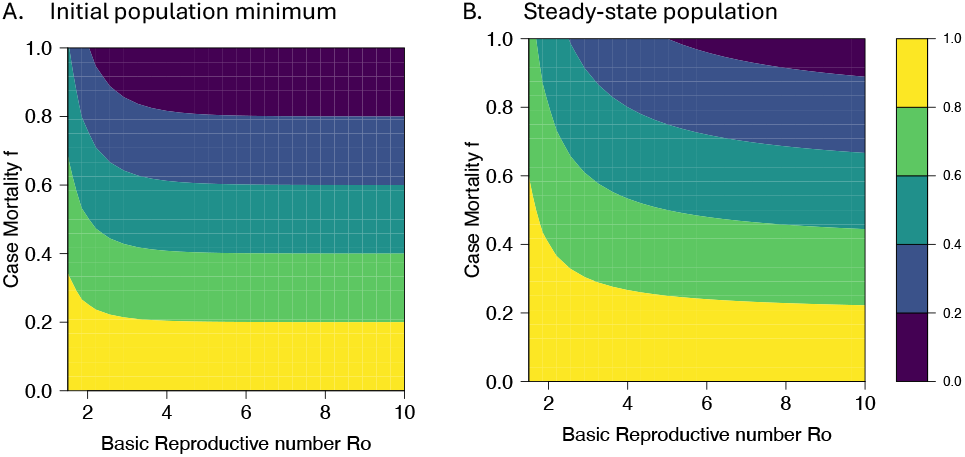
Initial vs long-term dynamics for a model with constant births and a fixed death rate. Outcome for initial dynamics on left using the final size formula and the long-term dynamics using the steady-state equilibrium. The population experiences modest recovery in the long-term.

#### Logistic growth

For this model, we assume that the host population exhibits logistic growth in the absence of the pathogen. Logistic growth is modeled by density dependence of both the birth and death rates. Here we assume that the birth rate decreases linearly with density [*b*(*N*) = *b*_0_ − *k*_*b*_*N*], and the death rate increases linearly with density [*d*(*N*) = *d*_0_ + *k*_*d*_*N*]. Through appropriate choice of parameter values, the carrying capacity (equilibrium population density) in the absence of pathogen is here always scaled to 1. The endemic equilibrium appears to be locally stable for the suite of parameter values of interest – at which the equilibrium exists (Supplement S2). In contrast to the previous model, homeostasis here operates through a combination of birth rates decreasing and death rates increasing when *N* > 1 and the reverse when *N* < 1.

As a biological reference, we use parameter values to represent a small mammal, such as a rabbit, which enables us to relate our results to the well documented example of Australian rabbits. We consider two scenario’s for the regulation of the host population in the absence of the pathogen. In the first scenario, the population exhibits rapid regrowth when culled, which we refer to as *rapid homeostasis*. Rapid homeostasis is achieved by choosing high values for *k*_*b*_ and *k*_*d*_, the density-dependent components of the birth and death rates. Slow homeostasis is achieved with smaller *k*_*b*_ and *k*_*d*_ values. Within these constraints, the pathogen-free equilibrium is maintained at 1 by appropriate choice of the four birth/death values. The parameter values are given in Table 1.

Fig 2 illustrates the impact of a pathogen on initial and long-term suppression of the host population, specifically comparing effects of rapid and slow homeostasis (Fig 2, top and bottom, respectively). The first column shows the dynamics of the host population that illustrate rapid and slow homeostasis. The second and third columns show the host population size at the initial minima and at steady-state, respectively, which indicate the extent to which the pathogen suppresses the population in the short term and long-term respectively. The fourth column shows the ratio of the population suppression in the steady-state (endemic) to the initial (epidemic) phase. Major effects of homeostasis are evident in all columns except the second – the initial minima. This insensitivity of initial minima to homeostasis is fully compatible with the assumption that the final size of the initial epidemic phase is insensitive to birth and death.

**Figure 2:**
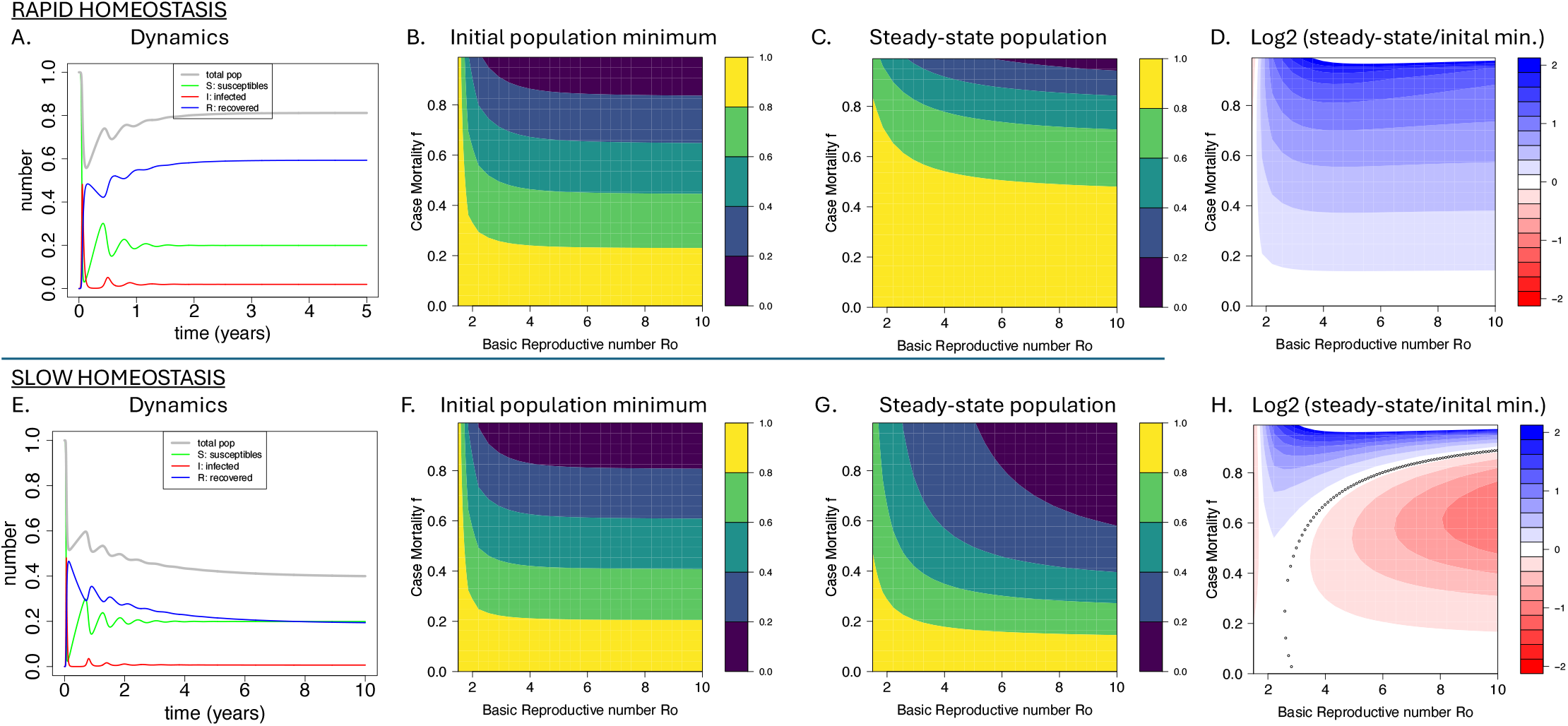
Population suppression following introduction of a pathogen. The top row shows the outcome when the homeostatic regulation of the host population is rapid, the bottom row for slow homeostasis. In the first column (panels A and E) shows the dynamics following introduction of a pathogen with an *R*_0_ = 5 and case mortality *f* = 0.5. We see that the initial suppression is similar regardless of the strength of homeostasis, but the population rebounds when homeostasis is rapid (panel A), whereas it declines further when homeostasis is slow (panel E). Panels B-C and E-F show how the initial and steady-state population suppression depend on *R*_0_ and *f*. We see similar initial population suppression for both rapid and slow homeostasis, but very different outcomes when steady-state is reached. The ratio of the population suppression at endemicity compared with during the epidemic phase for rapid and slow homeostasis is shown in Panels D and H. When there is rapid homeostasis the population invariably exhibits a rebound, and the steady-state population (panel C) is higher that at the initial decline (panel B). In contrast when homeostasis is slow, the population often declines further as steady-state is reached (panels F-H). The black points in panel H demarcate the separation between these two regimes as described by eqn 8. Parameter values are given in Table 1.

Panels A and E offer representative dynamics following the introduction of a pathogen with an *R*_0_ = 5 and *f* = 0.5 into a host population at its disease free equilibrium (*N* = 1). With rapid homeostatic regulation, the numbers rebound to a steady-state level that is significantly higher than the initial suppression – as is seen in the single trial of Fig 2A and by noting that the populations at steady state in 2C are higher than the initial minima in 2B. In contrast, if the host population experiences slow homeostasis, the long term population often declines beyond its initial minimum – as seen for a single trial in Fig 2E and by comparing panels 1F and 1G. A decline beyond the initial minimum does not apply throughout the parameter space, however, and panel H as well as a comparison of panels F and G show specific regions in which the final population size is larger or smaller than the initial minimum.

Further insight to this bifurcation behavior can be obtained with an analytical solution to an extreme case (Supplement S3). Assuming fast-recovery (*ν → ∞*), the population increases from its epidemic minimum if and only if

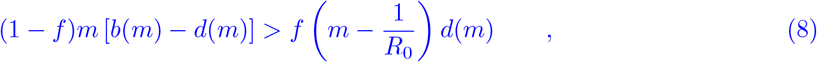

where *m* = 1 − *f F* is our approximation for the minimum size during the epidemic (*N*_*min*_). This result shows that the bifurcation has a general interpretation based on case mortality, *R*_0_, and the birth and death functions; it is not an anomaly of our choice of highly specific parameter values for the birth and death functions. As *m* involves a transcendental equation, it’s value must be calculated numerically. The LHS is the net population growth from the epidemic minimum times the surviving fraction if everyone was infected; it clearly shows that the net growth rate of the population (at the minimum) is critical to recovery. The RHS is the product of the death rate at the minimum times the case mortality times the extent to which the post-epidemic population is above the disease-control threshold of 1*/R*_0_. Setting eqn 8 to an equality, the equation demarcates the boundary in Fig. 2H between the regions where the population suppression at steady state is greater than the initial level of suppression during the initial epidemic.

Why does homeostasis have such a profound effect on the population size at the steady state? Inequality (8) doesn’t necessarily provide suitable intuition. With rapid homeostasis, following the initial suppression of the host population by the pathogen, the birth rate greatly exceeds the death rate, and this excess of births enables population rebound. The more interesting case is slow homeostasis, because the population continues an overall decline after the initial invasion. This case is most easily understood by considering the extreme of no homeostasis (i.e., *k*_*b*_ = *k*_*d*_ = 0, which requires *b*_0_ = *d*_0_ > 0). After pathogen invasion and the population has reached its early minimum, there will be no subsequent upward population growth because *b*_0_ = *d*_0_. In the short term, pathogen abundance is held in check from a combination of low host density and recovered individuals that cannot be reinfected. However, the ongoing birth and death will gradually replace recovered individuals with susceptible individuals. As susceptibles increase, despite no population growth, further outbreaks can occur, and the population declines further. Eventually, the population reaches a low point from which the pathogen cannot expand. The process is shown in Supplement S1 Fig 4, for extreme case in which no homeostasis exists. In less extreme cases, slow homeostasis merely fails to enable enough upward growth to offset the continued pathogen expansion as recovered individuals are replaced by susceptibles.

**Figure 3:**
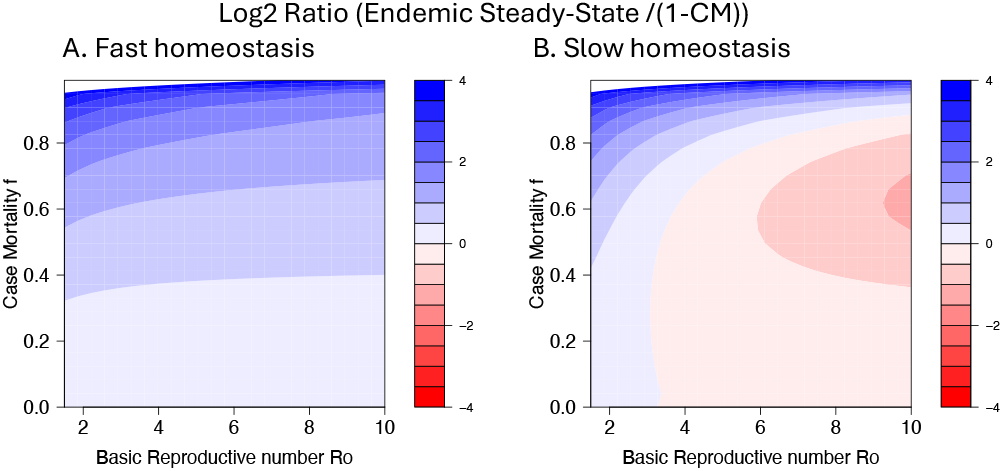
Population suppression at steady state can sometimes exceed the pre-introduction population times the case mortality, but not at the initial minimum. Red/pink shows the zones in which population size is smaller than 1 − *f*, which is the fraction of cases surviving infection. Thus, red and pink show the zones in which the population is suppressed beyond the level that would be imagined from experimental studies of case mortality. (A) is for fast homeostasis, (B) for slow homeostasis. Parameter values are given in Table 1.

**Figure 4:**
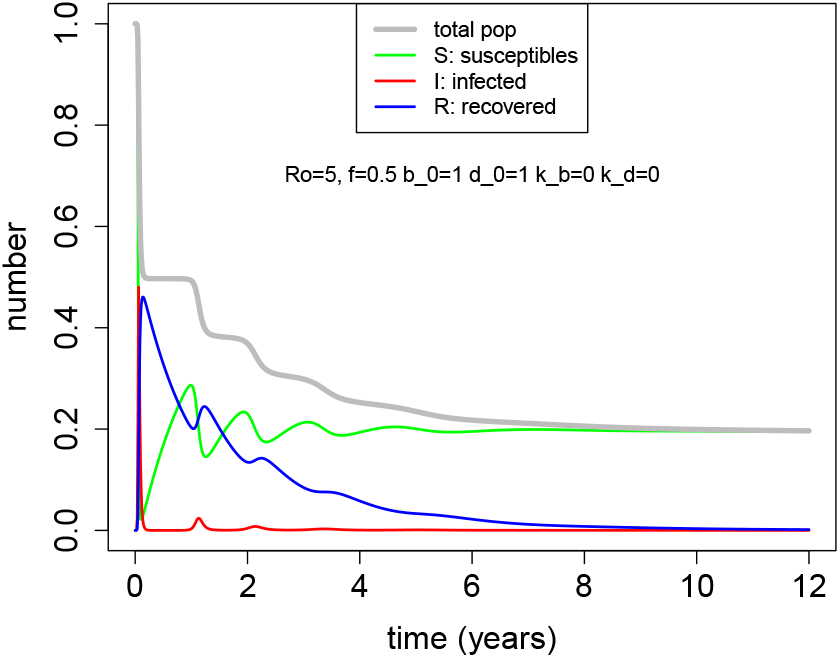
Pathogen invasion in the absence of homeostasis (*k*_*b*_ = *k*_*d*_ = 0). Following the initial invasion (the epidemic phase), the total population continues to decline until the reproductive number falls below 1. The early process involves stepwise decreases in population size, each preceded by an increase in the number of susceptibles that eventually enables the pathogen to expand further; recovereds decline as the susceptibles increase, so there is never a gain in total numbers. The decline ceases after the population falls to a fraction (1 − 1*/R*_0_) of the initial population, so the continued decline will happen provided the suppression during the initial epidemic phase does not exceed this this fraction.

#### Suppression can exceed case mortality

For extreme population suppression, an obvious criterion in the choice of pathogens is high case mortality (e.g., myxoma virus and calicivirus). But if the goal is population suppression, the question is whether high case mortality ensures high population suppression, or perhaps, whether high population suppression can be obtained without extremely high case mortality. Using the parameterizations of our logistic model in Table 1, figure 3 shows the relationship between *R*_0_, case mortality and endemic population size for rapid and slow homeostasis. Suppression exceeding case mortality can operate for our slow homeostasis values but not for our rapid homeostasis values. The effect is greatest for case mortality near *f* = 0.6.

#### Similar behavior for a second model of density dependence

The qualitative reasoning to explain the different trajectories following the initial population crash would suggest that the bifurcation is not unique to our linear model of density dependence. Supplement S4 shows that the same type of behavior is observed with a Beverton-Holt model of density dependence and further suggests that the linear model understates the generality of the bifurcation behavior.

## Discussion

This paper makes the simple point that, depending on the strength of density dependent population regulation, a pathogen that kills a fraction of the individuals it infects can cause its greatest population suppression either in the epidemic phase or in the endemic phase. Whereas the impact of the epidemic phase is largely independent of birth and death rates, the population size during the endemic phase depends fundamentally on birth and death rates. This finding complicates predicting the long term impact of novel, lethal pathogens. The results apply in the absence of any host or pathogen evolution, maternal antibodies, or age-dependence of infection severity, and inclusion of those factors would certainly modify these outcomes. So the results should be considered a baseline subject to many complications.

Our study builds on and integrates studies in a number of areas. The first is the role of density dependent regulation populations. Considerable effort by ecologists in the 1900s attempted to resolve whether populations were regulated by density-independent versus density-dependent mechanisms [4]. We have chosen a simple form of density dependence that gives rise to logistic growth and others are possible (one of which was included briefly). One might for example include models with an allee effect and stochasticity [23, 26]. We expect the allee effect to be equivalent to slower homeostasis when the population is low so we might expect that this would contribute to a further increase in the level of population suppression at steady state. We note that as the focus is on homeostatic regulation we do not consider a model lacking homeostasis in which the birth and death rates are constant and in which the host population grows exponentially at rate (*b* − *d*). A second area that our analysis contributes to is the analysis of the initial dynamics and final size of epidemics [3, 17, 21, 22]. Our work extends earlier work which calculate the final size (the total number of individuals infected) to consider the reduction in the host population in the first wave of infections following the introduction of a novel pathogen. The third area is the role of pathogens in regulating host populations [1, 2, 3, 8, 20]. These and other studies in this area indicate that the suppression of the host population by a pathogen is sensitive to many ecological factors, sometimes in unintutive ways [16]. In that broad sense, the dependence of long-term versus short-term suppression on density dependence should not be a surprise.

Our paper adds to the list of factors that dictate the extent to which a newly introduced pathogen can suppress a host population in the long term. Host and parasite evolution are others, although host evolution is likely to offset the suppression. Enumerating these factors and using simple models to understand their role in the suppression of host populations is a first step. A second step is to identify which factors might actually play a role, as might be done by comparing the models with well-documented historical studies that used, for example, myxoma and caliciviruses to control rabbit populations [5, 6, 9, 10, 19, 24, 27]. If the models appear consistent with the historical case studies, this could set the stage for the design of rational strategies that use pathogens as agents for the control of animal populations.

## Acknowledgments

We acknowledge the following support: JB: P20-GM104420; RA: U01 AI150747 and U01 AI144616. We thank Robert Holt, Bryan Grenfell, Viggo Andreassen and Andrew Dobson for discussion.

## Supplemental Information

### Use of AI tools

Portions of these supplemental works were carried out with the assistance of Claude (Anthropic) Opus 4.8 and Fable 5, May–July 2026. Specifically, the tool was used to assist with: (i) the analytical derivations reported here—the closed-form rebound/decline criterion and the reduced-Jacobian/Routh–Hurwitz stability analysis; (ii) the symbolic and numerical verification of those derivations (the `sympy` checks and the parameter-space eigenvalue sweep described in the supplement S2); (iii) the supporting simulation and analysis code; and (iv) drafting and typesetting of these notes. All AI-generated derivations, code, and text were reviewed, checked, and—where analytical—independently verified symbolically and numerically by the authors, who take full responsibility for the correctness and integrity of the results. The tool was not used to generate or interpret empirical data, and it is not an author of this work.

## File S1: Supplementary Information

### Parameter values

#### Estimating the parameters for small animal (rabbit)

We take these as reflecting rapid homeostasis.

1. From Google AI : 30 female offspring per year. Lifespan in the wild i.e., at equilibirium in the absence of the pathogen is about 1 year. We choose maximum lifespan in the wild of 5 years though in captivity they can live a bit longer.
2. Rescale carrying capacity c=1.
3. Maximum lifespan 5 years gives *d*_0_ = 1*/*5.
4. To get natural lifespan about 1 year *d*_0_ + *k*_*d*_ * *c* = 1 gives *k*_*d*_ = 1 − 1*/*5 = 0.8
5. 30 female offspring per year gives *b*_0_ = ln(30) ≈ 3.5
6. At steady-state (i.e. *c* = 1): (*b*_0_ − *k*_*b*_ * *c*) = (*d*_0_ + *k*_*d*_*c*) = 1 which gives *k*_*b*_ 2.5
7. We can maintain the same eqm death rate by *d*_0_ = 0.8 *k*_*d*_ = 0.2
8. We can maintain the same eqm birth rate by *b*_0_ = 1.5 *k*_*b*_ = 0.5

#### Parameters for slower homeostasis

1. We keep the same carrying capacity *c* = 1 2.
2. Let new *b*_0_ = 1.5 and *d*_0_ = 4*/*5 = 0.8
3. so new *k*_*d*_ = 1 − 4*/*5 = 1*/*5 = 0.2
4. and new *k*_*b*_ = 1.5 − 1 = 0.5

Note for simulations: During the initial epidemic phase the simulations often resulted in very low number of infecteds and populations becoming negative in simulations. We thus added a very small term equivalent to constant infections from outside of the order of 10^−5^ as can be seen in the code.

### Supplementary figure referenced in the text

## File S2: Local stability of the disease-present equilibrium

The question on the table is whether we can say anything analytic about the stability of the endemic equilibrium of eqn (1) — the fixed point with the disease present. The honest worry is that the equilibrium itself is ugly: solve for it in closed form and you get a radical expression that Mathematica reports but no one wants to read. The natural inference is that a stability analysis built on top of that root would be worse still, and so not worth attempting.

But the inference does not follow. Local stability never needs the explicit root: it needs the relations that define the fixed point, and those are clean. This note works the reduction through, lands the verdict, and reports a numerical search for any way to break it. The short version: the disease-present equilibrium is locally asymptotically stable everywhere we can make it exist, the analysis is a textbook cubic, and the horrendous radical never appears.

### The model and the three relations

We use eqn (1) with logistic homeostasis, *b*(*N*) = *b*_0_ − *k*_*b*_*N* and *d*(*N*) = *d*_0_ + *k*_*d*_*N*, transmission *β* = *R*_0_*ν*:

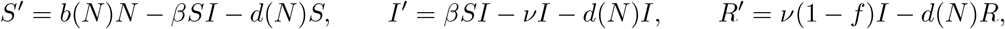

with *N* = *S* + *I* + *R*. One convention to flag at the outset: we write transmission as *β* = *R*_0_*ν*, the large-*ν* (fast-recovery) approximation the preprint already uses. The exact relation at the normalized disease-free state *S*_0_ = *N*_0_ = 1 is *β* = *R*_0_(*ν* + *d*_0_), since *R*_0_ = *β/*(*ν* + *d*_0_) there, so *β* = *R*_0_*ν* drops *d*_0_ against *ν*. Nothing in the reduction or the sweep below takes *ν → ∞* — both are carried out at finite *ν* — and this approximation’s only consequence, a shift in the meaning of the *R*_0_ axis at small *ν*, is quantified with the sweep. This is a three-dimensional system — births and deaths make *N* dynamic, so there is no conservation law to drop the dimension — and its Jacobian is therefore 3 × 3 and its characteristic polynomial a cubic. That single fact is most of the answer. A cubic has a Routh–Hurwitz criterion of three inequalities, and those inequalities are all we need.

At a disease-present fixed point (*I*\* > 0) the equations give three relations:

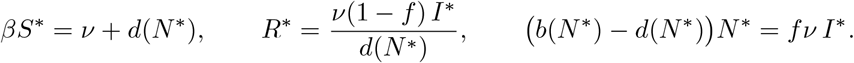

The first comes from *I*′ = 0, the second from *R*′ = 0, the third from *N* ′ = *S*′ + *I*′ + *R*′ = 0. None of them is the radical. They pin *S*\* and *R*\* to *N* * and *I*\*, and the closure *S*\* + *I*\* + *R*\* = *N* * leaves a single scalar equation for *N*\* — the quadratic in the *ν* limit, the transcendental at finite *ν*, both already in `criterion.py`. The stability analysis below uses the finite-*ν* transcendental, not the *ν → ∞* quadratic; the reduction that follows is exact at finite *ν*. We keep *N* * implicit: every coefficient below is a polynomial in the equilibrium coordinates, and any sign analysis reduces modulo *N* *‘s defining equation rather than substituting its root.

### The reduced Jacobian

Differentiate, then impose only *βS*\* = *ν* + *d*\* (writing *d*\* = *d*(*N* *) and 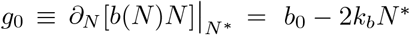, the marginal recruitment rate). The Jacobian at the endemic point is

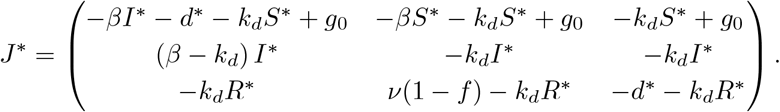

The structure is readable. The shared −*k*_*d*_*S*\*, −*k*_*d*_*I*\*, −*k*_*d*_*R*\* down each row is density-dependent mortality, *d*′(*N* *) = *k*_*d*_, acting on each compartment; the recruitment term *g*_0_ sits across the susceptible row because all births enter *S*. One entry deserves a flag, because it corrects a shorthand from our conversation: the (*I, I*) entry is **not** zero. The product rule on −*d*(*N*)*I* leaves a surviving piece, so

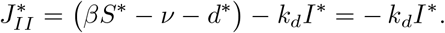

The first bracket vanishes by the *I*′ = 0 relation; the −*k*_*d*_*I*\* does not. It is zero only when death is density-independent (*k*_*d*_ = 0). This does not touch the cubic argument, but it should be written correctly.

### The cubic and the criterion

Write the characteristic polynomial as *λ*^3^ + *a*_1_*λ*^2^ + *a*_2_*λ* + *a*_3_, with *a*_1_ = − tr *J*\*, *a*_2_ the sum of the three principal 2 × 2 minors, and *a*_3_ = −det *J*\*. These are short. The trace term collapses to a clean closed form,

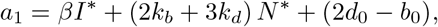

a linear expression in (*I*\*, *N* *) — positive across our parameter sets because *βI*\* and the demographic term dominate the constant 2*d*_0_ − *b*_0_ (which is itself negative for the slow envelope). The minor sum *a*_2_ is a degree-two polynomial in the equilibrium coordinates, and the determinant factors as

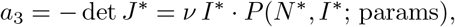

with *P* a polynomial. The explicit forms are in `coeffs_for_note.py`; the point for present purposes is that they are polynomials, not radicals. The Routh–Hurwitz criterion for a cubic then reads

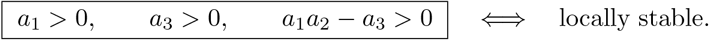

The factored determinant repays a second look. Because *a*_3_ ∝ *I*\*, the constant coefficient vanishes linearly as *I*\* → 0 — that is, as the endemic branch meets the disease-free equilibrium at the transcritical point. One root of the cubic crosses zero there, which is exactly the stability exchange we expect when the disease-free equilibrium loses stability and hands it to the endemic branch. The endemic point inherits its stability from that crossing, and nothing in the interior of the feasible region undoes it.

### Verdict, and an attempt to break it

We evaluated the cubic and its eigenvalues at the canonical sets. Every case is stable, with the same eigenstructure throughout: one negative real eigenvalue — the slow demographic recovery mode, the rate at which *N* relaxes back toward its endemic level — and a complex-conjugate pair with negative real part, the damped epidemic oscillation whose frequency scales with *ν*.

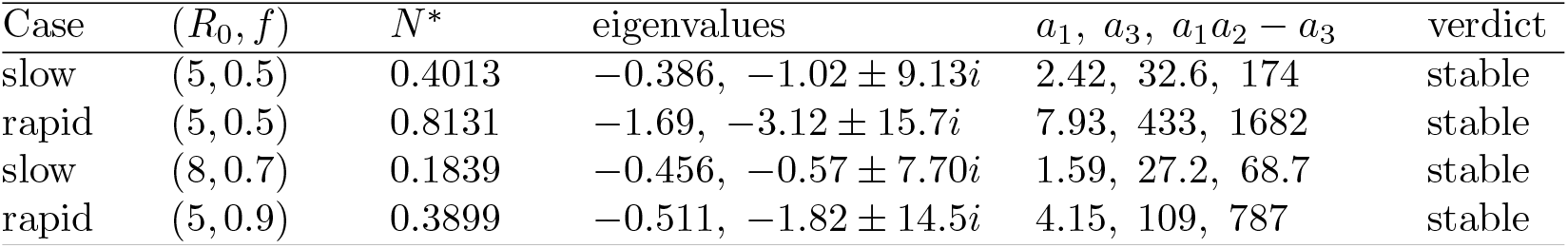

That the canonical points are stable is suggestive but not decisive, so we tried to break it. An independent sweep solved the endemic equilibrium and computed its eigenvalues over roughly 1.3 × 10^6^ feasible parameter sets: *R*_0_ from 1.05 to 30, *f* from 0.01 to 0.99, *ν* from 1 to 500, and the demographic rates *b*_0_, *d*_0_, *k*_*b*_, *k*_*d*_ over 0.01 to 10, both for the linear homeostasis above and for the Beverton–Holt form of S4. Not one feasible endemic equilibrium was unstable, and no eigenvalue pair crossed into the right half-plane — no Hopf bifurcation anywhere in the search. Because the sweep runs *ν* down to 1, the verdict is not a fast-recovery artifact. The *β* = *R*_0_*ν* convention leaves one caveat, and it is about labeling rather than stability: the swept *R*_0_ is the large-*ν* value, so the exact reproduction number is *R*_0_*/*(1 + *d*_0_*/ν*) — within about 2% of the label at *ν* = 50 but a factor of roughly 1.8 below it at *ν* = 1. That relabels the *R*_0_ axis at small *ν* without moving any eigenvalue across the imaginary axis, since the eigenvalues depend on the pair (*β, ν*) and not on how it is named. The least-stable case found had a dominant real part of −2.8 × 10^−4^, and it sat exactly where the theory says it should: at *I*\* → 0, the transcritical edge where *a*_3_ → 0, or in the limit of vanishingly small demographic rates where the slow mode’s timescale runs off to infinity. Scaling all rates by a factor *s* scales that eigenvalue by *s* without ever flipping its sign.

To be fair, a finite numerical sweep is not a proof of global-in-parameter stability, and we have not ruled out a Hopf bifurcation in some corner we did not sample. But the structure makes one unlikely: the cubic’s coefficients are positive and well-separated in magnitude across the whole feasible region, and *a*_1_*a*_2_ − *a*_3_ never approaches zero except at the transcritical edge, where it is *a*_3_ that vanishes, not the Hopf combination. The same reduction carries over unchanged to the Beverton–Holt homeostasis of S4 — only *b*′(*N*\*) and *d*′(*N*\*) change.

### What this buys the paper

The practical answer to the worry is the cleanest part. The radical equilibrium is an artifact of asking for the root; a stability analysis never asks for it. Substitute the relations into the Jacobian, read off a cubic, apply Routh–Hurwitz: the whole thing is a page of standard work, not a mess, and it is cheap to do either by hand on the reduced *J* * above or with a few lines of a computer-algebra system. So we should do it.

And it sharpens the rebound/decline story rather than complicating it. There is a single endemic equilibrium, it is locally stable throughout the biologically relevant region, and the approach to it is a damped oscillation — no limit cycles, no sustained endemic oscillation, no second attractor competing for the trajectory. The rebound-versus-decline question is therefore not a question about stability or about which attractor wins. It is a question about where the one stable endemic level sits: above the post-epidemic minimum *m*, and the population rebounds to it; below *m*, and the same stable point is reached by continued decline.

### Files

- `stability_probe.py` — reduced Jacobian, cubic, and eigenvalues at the canonical sets.
- `coeffs_for_note.py` — the reduced *J*\* entries and the explicit *a*_1_, *a*_2_, *a*_3_.
- `verify_endemic_stability.py` — independent re-derivation and the 1.3 × 10^6^-point adversarial sweep (linear + Beverton–Holt); full log in `endemic-stability-verification-2026-06-13.txt`.

## File S3: Supplemental Information: An analytical rebound/decline criterion for the SIR-with-homeostasis model

### The criterion

We work with the model in equation (1) of the preprint, with logistic homeostasis *b*(*N*) = *b*_0_ − *k*_*b*_*N, d*(*N*) = *d*_0_ + *k*_*d*_*N*, and the disease-free equilibrium normalized to *N* = 1 (so *b*_0_ − *d*_0_ = *k*_*b*_ + *k*_*d*_). Setting *a* = *b*_0_ − *d*_0_ and *c* = *k*_*b*_ + *k*_*d*_, the three equilibrium conditions are *S*\* = (*ν* + *d*(*N*\*))*/β* from *I*′ = 0, *R*\* = *ν*(1 − *f*)*I*\**/d*(*N* *) from *R*′ = 0, and (*a* − *cN* *)*N*\* = *fνI*\* from the sum (i.e., from *N* ′ = 0).

We can substitute these into the conservation constraint *S*\* + *I*\* + *R*\* = *N* *, multiply through, and collect powers of *N* *. The *I*\* term and the *d/ν* correction to *S*\* both vanish as *ν* → ∞, leaving

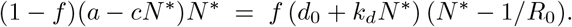

This is the simplification we wanted: a quadratic in *N* *. The positive root is

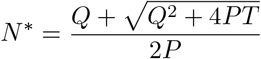

where

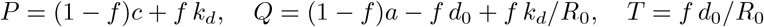

So far so good. The rebound criterion is *N* * > *m*, where *m* = 1 − *fF* (*R*_0_) is the post-epidemic minimum and *F* is the SIR final size *F* = 1 − exp(−*R*_0_*F*). After substituting and factoring, the criterion at the boundary becomes

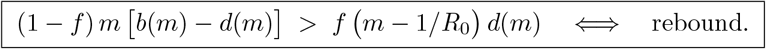

This is the result. The left side is the survival-weighted recovery rate at *N* = *m* — the rate at which the population would grow back toward *N* = 1 if disease deaths counted only with the survival fraction (1 − *f*). The right side is the disease drag — the rate at which the disease pulls the population down, weighted by the case mortality *f* and proportional to the susceptibility excess *m* − 1*/R*_0_ (how far above the disease-control threshold 1*/R*_0_ the post-epidemic population sits).

#### From N* to the criterion

The step from the radical to the boxed inequality looks worse than it is, and the reason is that the radical is a detour: the boxed inequality is nothing other than the quadratic itself, evaluated at m. Three observations get us there.

First, name the quadratic. Move everything in the fast-epidemic equilibrium relation to one side and define

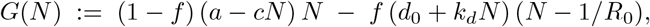

so that the equilibrium condition is exactly G(N*) = 0. Expanding and collecting powers of N gives *G*(*N*) −− *PN* ^2^ + *QN* + *T* with precisely the P, Q, T of the displayed root, which is the sense in which N* is “the positive root”: the quadratic *PN* ^2^ − *QN* − *T* = 0 is G(N) = 0 with the sign flipped.

Second, read G in terms of b and d rather than a and c. The normalization was *a* = *b*_0_ − *d*_0_ and *c* = *k*_*b*_ + *k*_*d*_, so *a* − *cN* = (*b*_0_ − *k*_*b*_*N*) − (*d*_0_ + *k*_*d*_*N*) = *b*(*N*) − *d*(*N*), and *d*_0_ + *k*_*d*_*N*) is just d(N). Substituting these two identities into the definition of G, with no rearrangement at all,

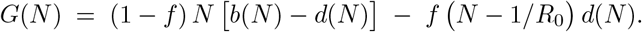

The right-hand side is the boxed criterion’s left side minus its right side, evaluated at N. So the boxed inequality is the statement G(m) > 0. Nothing has been substituted or factored; the two expressions were the same object written in two alphabets.

Third, and this is the only place any work happens, get from *G*(*m*) > 0 back to N* > m. Here the sign structure does everything. *P* = (1−*f*)*c*+*fk*_*d*_ > 0 and *T* = *fd*_0_*/R*_0_ > 0 for any f in (0,1), so G is a downward parabola with G(0) = T > 0. A downward parabola that is positive at the origin has one root on each side of it, which we can see directly from the radical: 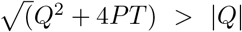 whenever PT > 0, so the two roots

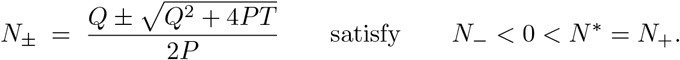

G is positive strictly between its roots, so for any positive N we have *G*(*N*) > 0 iff *N* < *N* *, the *N_*root being unreachable. And *m* = 1 − *f F* (*R*_0_) is positive, since *f* < 1 and *F* < 1 give *f F* < 1. Evaluating at N = m:

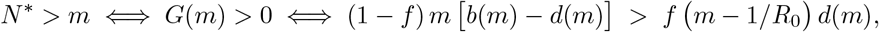

which is the boxed criterion. The radical never appears, which is the point: we do not need to know where N* is to know which side of m it falls on, only that G tips negative as N passes it.

For anyone who prefers to push the radical through, the same conclusion survives the squaring, though it costs a case split. N* > m reads 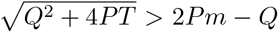. If the right side is negative the inequality holds automatically, and it holds consistently, since 2Pm < Q then forces *Gm* = −*Pm*^2^ + *Qm* + *t* > −*Pm*^2^ + 2*Pm*^2^ + *T* = *Pm*^2^ + *T* > 0. If the right side is non-negative, squaring is faithful and gives *Q*^2^*P* + 4*PT* > 4*P* ^2^*m*^2^ −4*PQm* + *Q*^2^. The *Q*^2^ cancels, and dividing by 4P > 0 leaves *T* > *Pm*^2^ − *Qm*, which is G(m) > 0 again. Both branches land on the same place, which is what the direct argument told us without the branching.

Pushing the f in the first term through the bracket gives the equivalent form quoted below: *G*(*m*) = (1−*f*) *m b*(*m*) −*m d*(*m*)[(1−*f*)+*f*] +*f/R*_0_ *d*(*m*) = *m*[(1−*f*) *b*(*m*) −*d*(*m*)]+(*f/R*_0_) *d*(*m*). Readers should be careful about one thing here. What the chain above establishes is an algebraic equivalence, N* > m iff the boxed inequality, and that is all it establishes. Whether N* > m is itself the right reading of “rebound” is a dynamical question rather than an algebraic one, since it presumes the endemic equilibrium is the state the post-epidemic population actually goes to. That presumption is not free, but it is now discharged separately: another Supplement section verifies that the endemic equilibrium is locally stable across the parameter region we work in.

The algebra here is verified symbolically in `verify_criterion_steps.py`, which confirms the *G* = −*PN* ^2^ + *QN* + *T* identity, the b/d identification, that both N ± are roots, the equivalent form, and the sign equivalence N* > m iff G(m) > 0 over 20,000 random parameter draws.

A clean equivalent form pushes f into the LHS bracket: rebound iff *m* · [(1 − *f*)*b*(*m*) − *d*(*m*)] + (*f/R*_0_) *d*(*m*) > 0. The first term is m times a “survival-weighted net growth rate” — what the population growth rate would be if a fraction f of births died immediately. For our slow-homeostasis parameters this term is negative at N = m (the population is below the survival-weighted carrying capacity), and rebound is achieved only if the small (f/*R*_0_) d(m) compensation is large enough.

### Why this should make some sense

Roughly put, the criterion compares two recovery races. The first is the demographic process: how fast can births close the gap between the epidemic minimum m and the disease-free equilibrium N = 1. The second is the endemic-disease process: once susceptibles re-accumulate past 1/*R*_0_, the disease starts taking new hosts at a rate proportional to d(m) and case mortality f. Whether the population settles above or below m depends on which of these processes is faster.

Two limits make the geometry transparent. When the homeostasis is shaped so that the survival-weighted carrying capacity (the root of (1 - f) b(N) = d(N)) lies above m, the demographic process wins and the system rebounds. For our slow parameters (*b*_*d*_ = 1.5, *d*_0_ = 0.8, *k*_*b*_ = 0.5, *k*_*d*_ = 0.2) at f = 0.5 we have (1−*f*)*b*_0_ − *d*_0_ = −0.05 < 0, so the survival-weighted recovery rate is negative for all N > 0 — the population cannot sustain itself if everyone catches the disease — and the only thing pulling N up from below is the f/*R*_0_ d(m) term. That term is small at high *R*_0_, which is why slow homeostasis tips into continued decline as *R*_0_ grows.

For our rapid parameters (b0 = 3.5, d0 = 0.2, kb = 2.5, kd = 0.8), the same calculation gives (1 − *f*)*b*_0_ − *d*_0_ = +1.55: the population is robustly able to absorb a 50% mortality event, and N* sits near the disease-free equilibrium across the entire (*R*_0_, f) plane we tested.

### Validation against simulation

We compared the closed-form *ν →* ∞ N* and a numerical root of the full transcendental S* + I* + R* = N* equation (i.e., keeping the I* and *d/ν* terms) against long-time integration of the deterministic ODE on a grid of (*R*_0_, f). The full set of comparisons lives in `validate_criterion.py`; a representative slice at our nominal *ν* = 50:

| Regime | $(R_0, f)$ | $N_{\min}$ | $N^*$ (sim) | $N^*$ (finite $\nu$) | $N^*$ ($\nu \rightarrow \infty$) |
| --- | --- | --- | --- | --- | --- |
| Slow | (5, 0.5) | 0.5035 | 0.4013 | 0.4013 | 0.3896 |
| Slow | (3, 0.5) | 0.5298 | 0.5380 | 0.5380 | 0.5261 |
| Slow | (8, 0.8) | 0.2003 | 0.1573 | 0.1573 | 0.1522 |
| Rapid | (5, 0.5) | 0.5035 | 0.8131 | 0.8131 | 0.8072 |
| Rapid | (8, 0.8) | 0.2003 | 0.4937 | 0.4937 | 0.4783 |

The finite-*ν* analytical N* agrees with simulation to four decimal places across the entire grid we tested. The *ν → ∞* closed form is within 1–3% of simulation; the discrepancy is the order-(*d/ν*) correction to S* and the order-(1/*ν*) contribution from I*, both of which are small at *ν* = 50 and vanish in the fast-epidemic limit.

The corresponding picture in (*R*_0_, f) is in Figure 1. The black solid line is the *ν → ∞* boundary; the black dashed line is the boundary computed from the finite-*ν* transcendental. They differ by about 0.03 in f. Simulation outcomes (open blue circles for rebound, red crosses for continued decline) sort cleanly across the boundary in both panels.

**Figure 1:**
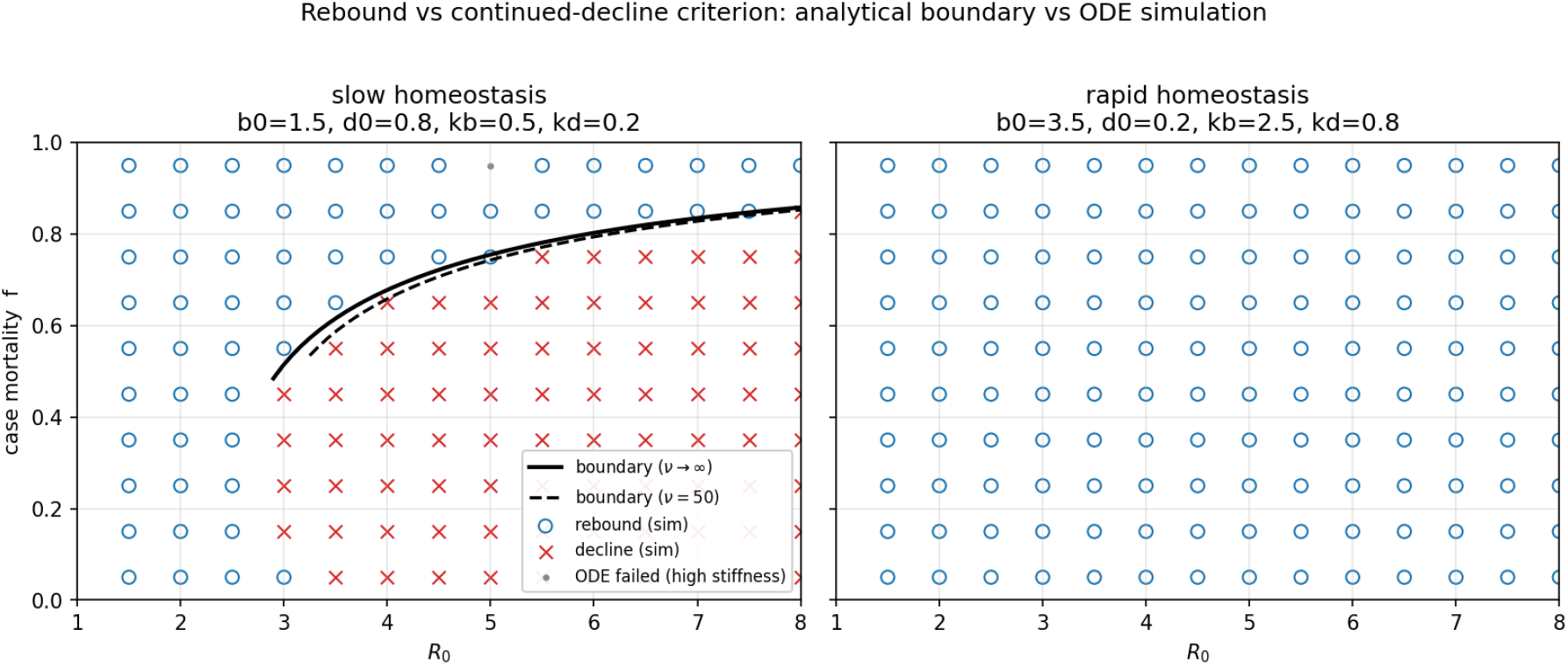
Rebound/decline boundary vs simulation, slow and rapid homeostasis at *ν* = 50, on a 14 × 10 (*R*_0_, f) grid. The solid curve is the *ν* → ∞ closed-form boundary; the dashed curve is the boundary from the finite-*ν* transcendental. The single gray point at (*R*_0_ = 5, f=0.95) in the slow panel marks an ODE-solver stiffness failure; both analytical methods predict rebound there, matching every adjacent simulation.

A few observations from the figure. First, the rapid-homeostasis panel is entirely rebound across the (*R*_0_, f) plane we tested. The (1 - f) *b*_0_ − *d*_0_ > 0 condition is robust here, and we would have to push f very close to 1 — beyond biological plausibility — to flip the verdict. Second, the slow-homeostasis decline region opens up around *R*_0_ ≈ 3 and grows with *R*_0_, but the upper boundary remains below f *approx* 0.85 even at *R*_0_ = 8: very high case mortality flips the verdict back to rebound because the epidemic itself drives N below the endemic floor 1*/R*_0_. Third, the *ν → ∞* and finite-*ν* boundaries are close enough that the closed form is adequate for most practical purposes; the finite-*ν* formula is available when an extra decimal point of precision matters.

To be fair, there is a small numerical caveat: at *R*_0_ ≳ 3 with *f* ≳ 0.9, the LSODA integrator hits stiffness limits and we fall back to Radau and then BDF. One point in our grid (*R*_0_ = 5, f = 0.95, slow homeostasis) defeated all three and we leave it as a missing point in the figure. The analytical N* (both *ν → ∞* and finite-*ν*) handles this regime cleanly, which is actually one argument for the analytical work: the simulator’s weakest regimes are exactly the ones where the criterion is most needed.

### Files

- `criterion.py` — closed-form *ν → ∞* N*, finite-*ν* numerical N*, criterion sign function, boundary solver.
- `validate_criterion.py` — point-by-point comparison and figure generation.
- `verify criterion steps.py` — symbolic (sympy) verification of the N* → criterion algebra, plus a randomized check of the sign equivalence.
- `sir_model.py` — the deterministic ODE and stochastic SSA; the integrator now falls back from LSODA → Radau → BDF on stiffness/overflow.
- `criterion-2026-05-12.png` — Figure 1 above.

## Supplement S4: Beverton-Holt

In the fast-epidemic limit the boundary between post-epidemic rebound and continued decline is the sign of a single quantity,

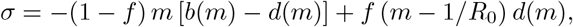

evaluated at the post-epidemic minimum *m* = 1 − *f F* (*R*_0_). We derived that criterion for linear (logistic) homeostasis, *b*(*N*) = *b*_0_−*k*_*b*_*N* and *d*(*N*) = *d*_0_+*k*_*d*_*N* . It reproduces the simulated boundary closely, and for slow homeostasis it draws a decline region that grows with *R*_0_.

But a criterion is only as general as the functional form it assumes. The decline region might be a real feature of density-dependent recovery, or it might be an artifact of having written density dependence as a straight line. So the question we take up here is not where the boundary moves when we change the form. It is whether the boundary is there at all.

The answer reframes the contribution. For slow homeostasis the boundary is robust: it survives the change of form, and saturating the mortality only enlarges it. The surprise is on the other side. For rapid homeostasis, which the linear criterion clears as certain to rebound, a realistic saturating mortality manufactures a decline region where linear theory saw none. Linear homeostasis, in other words, understates the risk.

### Sharpening the question

To ask whether a boundary exists, we have to hold something fixed. Otherwise the question is empty: with the intrinsic growth rate free we could make any model rebound everywhere by raising it, or decline everywhere by lowering it. So we hold the demographic envelope fixed and vary only the shape of the density dependence.

Concretely, we keep the intrinsic per-capita rates *b*(0) = *b*_0_ and *d*(0) = *d*_0_, and we keep the carrying capacity by pinning the disease-free equilibrium at *N* = 1. Within those constraints we replace the straight lines with a both-rates-saturating Beverton–Holt form,

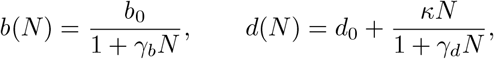

with *κ* fixed by the equilibrium condition *b*(1) = *d*(1). The two curvatures *γ*_*b*_ and *γ*_*d*_ are the only things that differ from the linear model. We put the saturating form on both rates, rather than on births alone as textbook Beverton–Holt does, because the preprint’s homeostasis acts on births and deaths together, and we want the comparison to keep that structure.

It is worth noting that the linear model is not a limiting case of this family. Setting *γ*_*b*_ = 0 gives constant births, not the linear decline *b*_0_ − *k*_*b*_*N* . So the comparison is between two genuinely different shapes at a shared envelope, not between a curve and its own small-curvature limit. One constraint falls out of the calibration and matters below: pinning the DFE requires *b*_0_*/*(1+*γ*_*b*_) > *d*_0_, so the birth-saturation cannot exceed *γ*_*b*_ < *b*_0_*/d*_0_ −1. For the slow envelope that ceiling is only 0.875.

### The rapid envelope understates its own risk

Under linear homeostasis the rapid envelope (*b*_0_ = 3.5, *d*_0_ = 0.2) rebounds across the entire (*R*_0_, *f*) plane we tested. There is no boundary. The first surprise is how easily one appears once the mortality saturates. Figure 1 (left) maps the fraction of the (*R*_0_, *f*) plane in continued decline over the curvature plane; the black region is where the population still rebounds everywhere, and the cyan curve marks the onset of a decline region.

**Figure 1:**
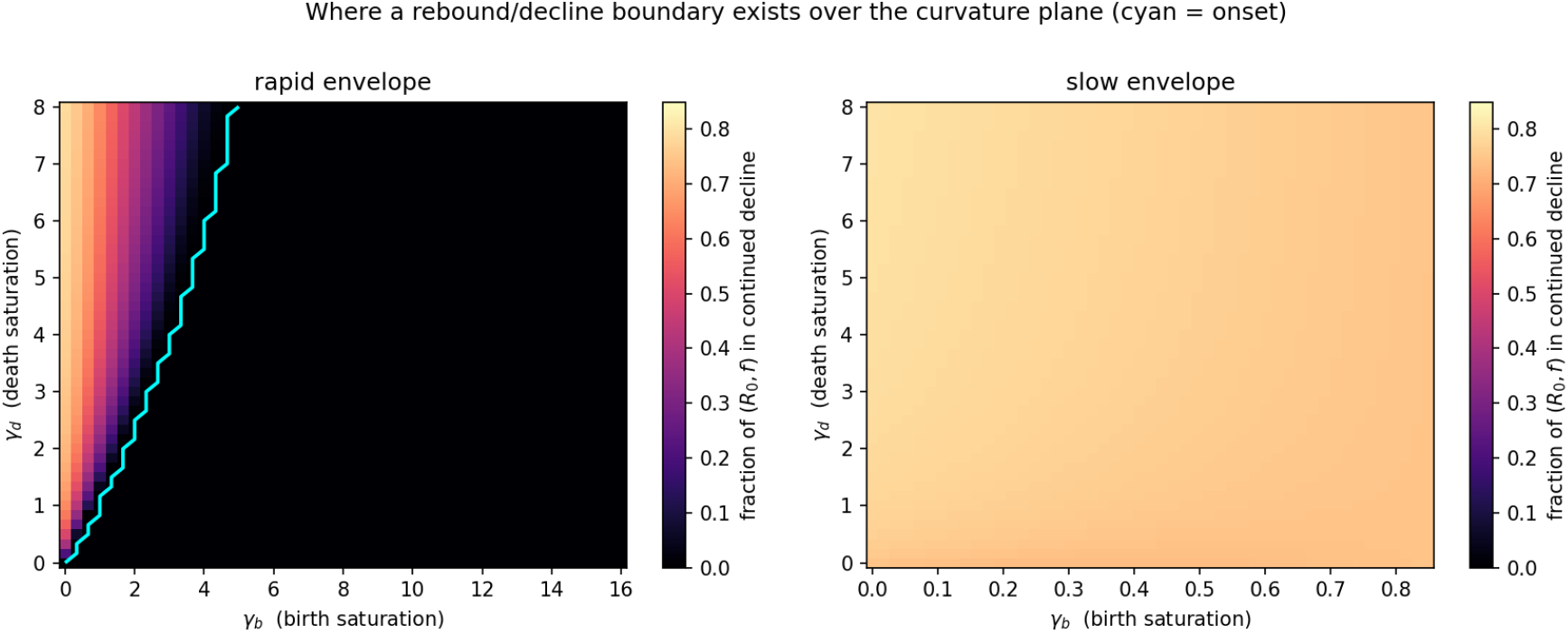
Fraction of the (*R*_0_, *f*) plane in continued decline, over the curvature plane (*γ*_*b*_, *γ*_*d*_), at fixed envelope. Left: rapid homeostasis, where a boundary exists only above the cyan onset curve. Right: slow homeostasis, where a decline region is present at every feasible curvature (note the truncated *γ*_*b*_ axis: birth-saturation is capped at 0.875 for this envelope).

Reading up the left edge of the rapid panel, at *γ*_*b*_ = 0 the decline fraction climbs from zero at *γ*_*d*_ = 0 to 64% at *γ*_*d*_ = 1, and to 79% by *γ*_*d*_ = 8. Death-saturation on its own turns a population that linear theory calls safe into one that declines across most of the epidemic-severity plane. Birth-saturation pulls the other way. Reading across at *γ*_*d*_ = 4, the decline fraction falls from 77% at *γ*_*b*_ = 0 to 23% at *γ*_*b*_ = 2, and back to zero by *γ*_*b*_ = 4. So the existence of a boundary is the outcome of a competition between the two curvatures.

Figure 2 (top row) shows the same story in the (*R*_0_, *f*) plane itself. At *γ*_*d*_ = 0 the plane is all rebound. Turn up the death-saturation and a decline cone opens from the high-mortality corner and widens. The open circles and crosses are independent ODE checks at finite *ν*; they sort across the analytical boundary as they should.

**Figure 2:**
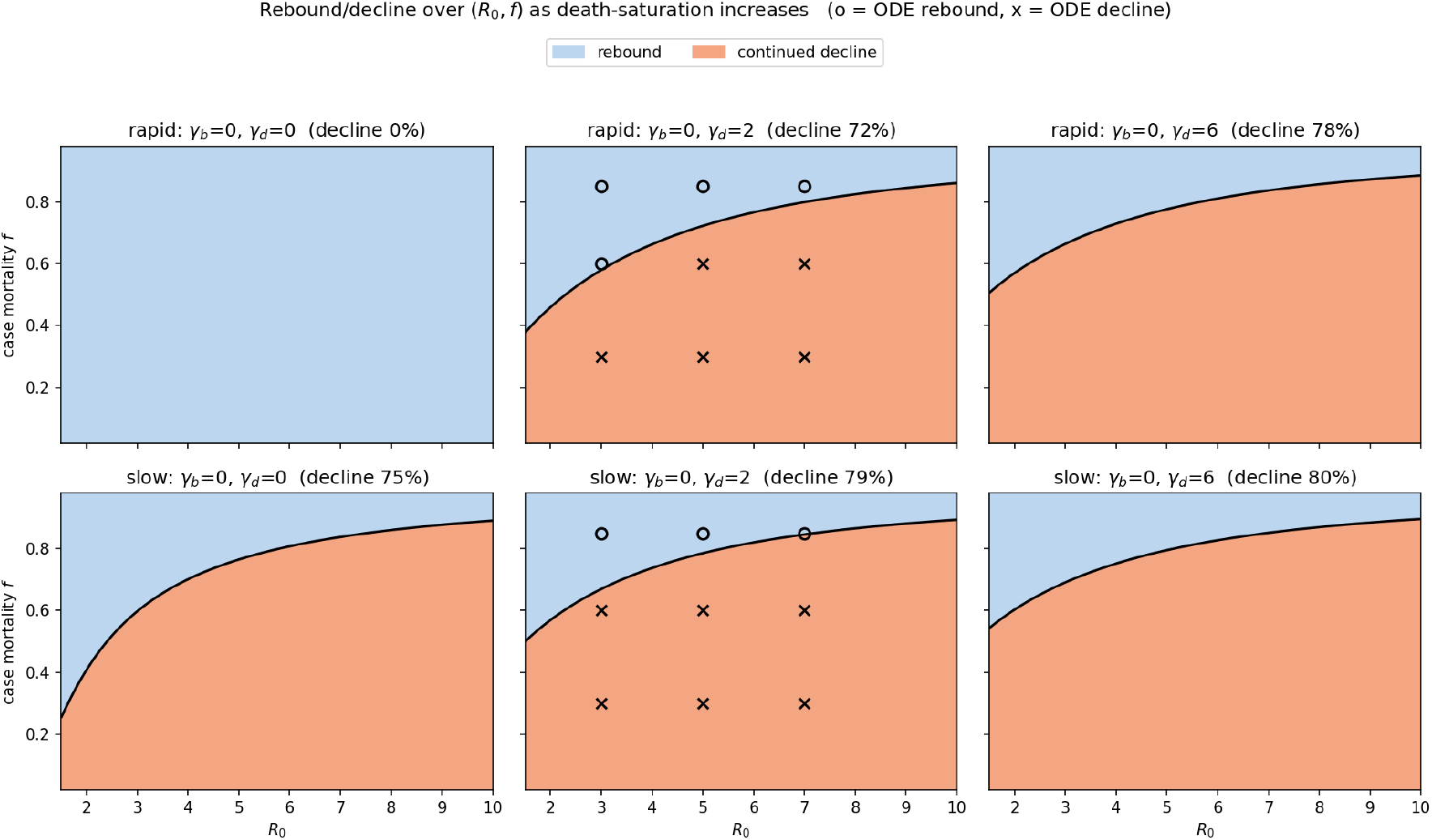
Rebound (blue) versus continued decline (orange) over (*R*_0_, *f*) as death-saturation *γ*_*d*_ increases, at *γ*_*b*_ = 0. Top: rapid envelope, where the decline cone is absent at *γ*_*d*_ = 0 and emerges with saturation. Bottom: slow envelope, where it is present throughout. Black curve is *σ* = 0; markers are finite-*ν* ODE outcomes (o = rebound, x = decline).

### The slow envelope: the decline region is not an artifact

The slow envelope (*b*_0_ = 1.5, *d*_0_ = 0.8) gives the cleaner answer to our original question. Figure 1 (right) is orange everywhere: a decline region exists at every curvature we can reach, and its size barely moves, from 75% of the plane at *γ*_*d*_ = 0 to 80% at *γ*_*d*_ = 8. The decline phenomenon the preprint identified is therefore not a consequence of writing homeostasis as a straight line. It survives the change of form, and saturating the mortality only enlarges it.

We say “every curvature we can reach” advisedly. The slow envelope’s birth-saturation is capped at *γ*_*b*_ < 0.875, so we cannot test whether very strong birth-saturation would rescue it the way it rescues the rapid envelope. Across the feasible range, though, the slow boundary is essentially immovable.

### Why this makes sense

Roughly put, the criterion is a race between the births that rebuild the population and the deaths, ordinary and disease-driven, that hold it down, both read at the post-epidemic minimum *m*. Saturating the two rates changes who wins, and it does so through their behavior at low density.

Consider the mortality first. A saturating death rate rises steeply with density at first and then plateaus. Pinning the DFE pushes that plateau up close to *b*(1) when *γ*_*d*_ is large, so the per-capita death rate stays near its ceiling even after the epidemic has thinned the population. Mortality fails to relax as the population falls, and the recovery stalls. That is exactly the regime in which the disease-drag term wins and the population keeps declining.

Births work the opposite way. A saturating birth rate climbs back toward its intrinsic value *b*_0_ as density drops, so a thinned population gets a strong per-capita birth boost, and the lower the density the harder it pushes back. Not surprisingly, then, birth-saturation favors rebound and death-saturation favors decline. The linear model sits at neither extreme, and for the rapid envelope it happens to sit on the safe side of a boundary that even mild death-saturation crosses. On reflection this makes the death side the one to watch. Mortality that does not ease at low density, whether from predation that is slow to switch to other prey or from a resource floor that bites regardless of density, is the realistic case rather than the exception.

### Validation

Three checks support the figures. First, the form-agnostic *σ* reproduces the closed form to six decimals on the linear model, so generalizing the criterion did not break the earlier result. Second, on a panel of twelve (*R*_0_, *f, γ*_*b*_, *γ*_*d*_) cells spanning both envelopes and both verdicts, the analytical sign of *σ* agrees with long-time integration of the full ODE at *ν* = 50 in all twelve, including the rapid decline cells that linear theory would not predict. Third, the endemic equilibrium stays unique: across the curvature and (*R*_0_, *f*) grids we searched, the *ν* → ∞ balance never had more than one feasible root. So the boundary is a simple separatrix rather than the edge of a fold, and the single-root logic behind *σ* holds throughout.

## To be fair

Several limits are worth stating plainly. The criterion is the fast-epidemic (*ν → ∞*) result, checked against finite *ν* at sample points rather than proven for finite *ν*. The both-rates-saturating form is a generalization of textbook Beverton–Holt, which saturates births alone, so a reader who reserves the name for the recruitment-only model would call ours a saturating-homeostasis variant, and that is fair. We held the mortality to forms that rise and plateau, not forms that accelerate, since saturation was the question; an accelerating mortality is a separate study. And the curvature ranges, though wide, are finite. None of these touches the qualitative result, but each marks a place where the present note stops.

### Let’s take stock

What the Beverton–Holt analysis adds is less a correction than a sharpening of where the risk lives. The slow-homeostasis decline the preprint identified is robust: it is a feature of density-dependent recovery, not of a linear approximation to it. The new part is on the other side. A population with fast homeostasis, which the linear criterion clears as certain to rebound, can be pushed into sustained decline by a mortality that saturates rather than rising without bound. To the extent that real density-dependent mortality plateaus, the linear model understates the risk of post-epidemic collapse.

This points the follow-up in a definite direction. The death-saturation curvature *γ*_*d*_ is the parameter that manufactures the danger, so the empirical question for any candidate system is how its mortality behaves at low density, not just what its intrinsic rates are. The natural next form is the *θ*-logistic, which interpolates between the linear and saturating shapes with a single exponent and would let us trace the onset of the boundary as a continuous function of curvature. There is more to do on the finite-*ν* correction as well. But the main lesson is already firm enough to carry: once homeostasis saturates, fast homeostasis is no longer a guarantee.

### Files

- `bh_existence.py` — curvature sweep, sign maps, validation table, and fold check; regenerates both figures.
- `criterion.py` — now carries the form-agnostic criterion `criterion_sigma` and the endemic-root counter `endemic_roots_inf_nu` alongside the linear closed form.
- `sir_model.py` — adds the Beverton–Holt forms behind `Params(form=“bh”)` and the DFE-pinned constructor `Params.bh_from_envelope`.
- `criterion-bh-curvature-2026-05-21.png` — Figure 1 (curvature existence map).
- `criterion-bh-signmaps-2026-05-21.png` — Figure 2 (*R*_0_, *f* sign maps as death-saturation increases).

## Notes

### Competing Interest Statement

The authors have declared no competing interest.

### Summary of Updates

We have addressed the following: 1. The stability of the host population at steady-state. 2. Identified the boundary between host population rebound vs further decline between initial minima and steady state.

### Funding

- NIH: P20-GM104420; U01 AI150747 and U01 AI144616

## References

[1] R. M. Anderson and R. M. May. Population biology of infectious diseases: Part i. Nature, 280 (5721):361–7, Aug 1979.

[2] V. Andreasen. Disease regulation of age-structured host populations. Theor Popul Biol, 36(2): 214–39, Oct 1989.

[3] V. Andreasen. The final size of an epidemic and its relation to the basic reproduction number. Bull Math Biol, 73(10):2305–21, Oct 2011.

[4] H. G. Andrewartha and C. Birch. The Distribution and Abundance of Animals. University of Chicago Press, 1954.

[5] B. Cooke, P. Chudleigh, S. Simpson, and G. Saunders. The economic benefits of the biological control of rabbits in Australia, 1950–2011. Australian Economic History Review, 53(1):91–107, 2013.

[6] T. Cox, T. Strive, G. Mutze, and W. Peter. Benefits of Rabbit biocontrol in Australia. Invasive Animals Co-operative Research Centre, Invasive Animals Co-operative Research Centre, Bruce, Canberra [Australian Capital Territory], 2013.

[7] A. P. Dobson and R. M. May. Patterns of invasions by pathogens and parasites. In H. A. Mooney and J. A. Drake, editors, Ecology of Biological Invasions of North America and Hawaii, pages 58–76. Springer, New York, New York, NY, 1986.

[8] C. S. Elton. Ecology of Invasions: By Animals and Plants. Methuen, London, 1958.

[9] F. Fenner and F. Ratcliffe. Myxomatosis. Cambridge university press, Cambridge, 1965.

[10] D. Fouchet, S. Marchandeau, M. Langlais, and D. Pontier. Waning of maternal immunity and the impact of diseases: the example of myxomatosis in natural rabbit populations. J Theor Biol, 242(1):81–9, Sep 2006.

[11] S. Gandon, M. J. Mackinnon, S. Nee, and A. F. Read. Imperfect vaccines and the evolution of pathogen virulence. Nature, 414(6865):751–6, Dec 2001.

[12] W. M. Getz and J. Pickering. Epidemic models: Thresholds and population regulation. The American Naturalist, 121(6):892–898, 1983.

[13] B. T. Grenfell and A. P. D. (editors). Ecology of infectious diseases in natural populations. Cambridge University Press, 1995.

[14] A. Handel, I. M. Longini, Jr, and R. Antia. What is the best control strategy for multiple infectious disease outbreaks? Proc Biol Sci, 274(1611):833–7, Mar 2007.

[15] M. E. Hochberg. The potential role of pathogens in biological control. Nature, 337(6204): 262–5, Jan 1989.

[16] R. D. Holt and M. Roy. Predation can increase the prevalence of infectious disease. Am Nat, 169(5):690–9, May 2007.

[17] W. O. Kermack and A. G. Mckendrick. Contributions to the mathematical-theory of epidemics .1. (reprinted from proceedings of the royal society, vol 115a, pg 700-721, 1927). Proceedings of the Royal Society A, 115:700–721, 1927.

[18] P. K. Kollepara, R. H. Chisholm, I. Z. Kiss, and J. C. Miller. Ethical dilemma arises from optimizing interventions for epidemics in heterogeneous populations. J R Soc Interface, 21 (211):20230612, Feb 2024.

[19] S. Marchandeau, D. Pontier, J.-S. Guitton, J. Letty, D. Fouchet, J. Aubineau, F. Berger, Y. Léonard, A. Roobrouck, J. Gelfi, B. Peralta, and S. Bertagnoli. Early infections by myxoma virus of young rabbits (oryctolagus cuniculus) protected by maternal antibodies activate their immune system and enhance herd immunity in wild populations. Vet Res, 45:26, Mar 2014.

[20] R. M. May and R. M. Anderson. Population biology of infectious diseases: Part ii. Nature, 280(5722):455–61, Aug 1979.

[21] J. C. Miller. A note on the derivation of epidemic final sizes. Bull Math Biol, 74(9):2125–41, Sep 2012.

[22] M. M. Nguyen, A. S. Freedman, S. A. Ozbay, and S. A. Levin. Fundamental bound on epidemic overshoot in the sir model. J R Soc Interface, 20(209):20230322, Dec 2023.

[23] E. P. Odum, G. W. Barrett, et al. Fundamentals of ecology. Saunders Philadelphia, 1971.

[24] G. Saunders, B. Cooke, K. McColl, R. Shine, and T. Peacock. Modern approaches for the biological control of vertebrate pests: an Australian perspective. Biological Control, 52(3): 288–295, 2010.

[25] M. E. Scott and A. Dobson. The role of parasites in regulating host abundance. Parasitol Today, 5(6):176–83, Jun 1989.

[26] P. A. Stephens, W. J. Sutherland, and R. P. Freckleton. What is the allee effect? Oikos, 87 (1):185–190, 1999.

[27] P. L. Taggart, T. W. O’Connor, B. Cooke, A. J. Read, P. D. Kirkland, E. Sawyers, P. West, and K. Patel. Good intentions with adverse outcomes when conservation and pest management guidelines are ignored: A case study in rabbit biocontrol. Conservation Science and Practice, 4(4):e12639, 2022.

